# TANGO2 deficient iPSC-differentiated cardiomyocyte and dermal fibroblasts have normal mitochondrial OXPHOS function

**DOI:** 10.1101/2022.06.27.497853

**Authors:** Weiyi Xu, Yingqiong Cao, Lorren Cantú, Eleni Nasiotis, Seema R. Lalani, Christina Y. Miyake, Lilei Zhang

## Abstract

Bi-allelic loss-of-function mutations in *TANGO2* (Transport and Golgi Organization protein 2) cause a rare multiorgan genetic disorder. Despite normal cardiac function at baseline, patients may experience lethal cardiac arrhythmias during “crises” often associated with metabolic stresses such as fasting, viral illness and fever. The molecular function of TANGO2 remains largely unknown. Previous studies have suggested a functional association with the mitochondrion, however definitive evidence is lacking. Further, functional impact of TANGO2 deficiency on mitochondrial function has not been investigated in a cardiac model. In this study, we utilized a recently developed patient-derived induced pluripotent stem cell differentiated cardiomyocytes (iPSC-CM) model by our group, along with patient-derived dermal fibroblast model, to interrogate whether loss of TANGO2 function leads to defective mitochondrial function. Both baseline and fasting condition were investigated. Oxygen consumption rate (OCR) was measured in Seahorse assays to assess mitochondrial function *in vitro*. The results showed both TANGO2 deficient dermal fibroblasts and iPSC-CM had no apparent defects in mitochondrial oxidative phosphorylation (OXPHOS) function under either baseline or fasting condition. Based on our study, we conclude that the lethal cardiac arrhythmias in TANGO2 patients are unlikely to be related to impaired mitochondrial OXPHOS function in the cardiomyocytes.

## 1 INTRODUCTION

TANGO2-related disease is a rare multiorgan genetic disorder caused by bi-allelic loss-of-function mutations in *TANGO2* (Transport and Golgi Organization protein 2) gene (1–14). Upon metabolic stresses such as fasting, dehydration, fever, and heat, TANGO2 patients often present with acute life-threatening metabolic crisis which may include hypoglycemia, hyperammonemia, rhabdomyolysis, and cardiac arrhythmias. Lethal cardiac arrhythmias during these crises are the leading cause of mortality in TANGO2 patients despite their normal cardiac function and rhythm at baseline (2, 3, 5, 7, 13, 15, 16). The lack of understanding in TANGO2 molecular function and the disease etiology greatly hinders the development of effective therapy for TANGO2-associated cardiac crises. As a result, current treatments are only supportive and with poor response.

The clinical resemblance of metabolic crises between TANGO2-related disease and inborn errors of metabolism especially those in the long chain fatty acid oxidation pathway has led to a speculation that a metabolic defect may be the primary driving force in the TANGO2-related disease associated crises. Moreover, the N-terminus of TANGO2 protein was predicted to consist of a mitochondrial targeting sequence (2), suggesting a potential function in the mitochondria. However, whether TANGO2 deficiency causes defective mitochondrial function remains largely controversial in the field (2, 5, 13). The discrepancy among different studies is likely attributed by individual variations in patient-derived dermal fibroblasts in which most of the experiments were carried out and the very limited sample size. Moreover, whether findings obtained from dermal fibroblast can be readily transferred to the heart is uncertain. To minimize the inter-individual variations, we have established multiple patient-derived induced pluripotent stem cell differentiated cardiomyocytes (iPSC-CM) model along with isogenic controls through ectopic expression of adenoviral vectors. This model successfully recapitulated key cardiac phenotypes seen in TANGO2-related disease, which supports the further use of these models to investigate the molecular mechanisms (ref). Here we take advantage of our existing iPSC-CM models, and generated patient-derived dermal fibroblasts and corresponding isogenic lines, to comprehensively interrogate the involvement of TANGO2 in mitochondrial OXPHOS function. Our data showed normal oxidative phosphorylation (OXPHOS) function in TANGO2 deficient fibroblast or iPSC-CM, suggesting primary defects in mitochondrial OXPHOS is unlikely to be the cause of TANGO2-associated cardiac crisis and arrhythmias.

## 2 MATERIALS AND METHODS

### 2.1 Cell culture

Primary dermal fibroblasts were isolated from a TANGO2 patients carrying exon 3-9 deletion mutation in *TANGO2* gene (ref). Cells were expanded and maintained in Dulbecco’s Modified Eagle’s Medium (DMEM) (GlonGen, 25-500) with 10% FBS (Thermo Fisher Scientific) in a humidified incubator at 37 degrees with 5% CO_2_. Fibroblasts with passage number between 4-10 were used for experiments. iPSC-CM was generated as previously described (ref) using GiWi method (17) or STEMdiff™ Ventricular Cardiomyocyte Differentiation Kit (STEMCELL Technologies, 05010), then further purified by lactate selection (18). iPSC-CM was maintained in RPMI1640 (Thermo Fisher Scientific, 11875119) supplemented with 1% B27 (Thermo Fisher Scientific, 17504001). Adenovirus expressing WT-TANGO2, G154R-TANGO2, and GFP control were generated by VectorBuilder as previously described (ref).

### 2.2 RNA extraction and reverse transcription

RNA was extracted using high pure RNA isolation kit (Roche, 11828665001). Reverse transcription was performed using iScript™ Reverse Transcription Supermix (Bio-Rad Laboratories, 1708841).

### 2.3 Genotyping

Primer pairs targeting exon 1-2 (Forward: 5’-CAGGCTGCTTGAAGACCTCG-3’; Reverse: 5’-ACGCGTTTTTGGAAACAGGG-3’) and exon 7-8 (Forward: 5’-CAACAATGAAGAGGCGCAGC-3’; Reverse: 5’-GTACTTGCTCAGCATGGGCTG-3’) were used to genotype the exon 3-9 deletion mutation. cDNA from fibroblasts carrying exon 3-9 deletion mutation and a WT control fibroblast were used as template for PCR. The PCR product was separated in a 2% agarose gel and imaged by ChemiDoc™ Touch Imaging System (Bio-rad Laboratories).

To sequence the WT-or G154R-TANGO2 expressed by adenoviral construct, a 747bp region containing the G154R mutation site was amplified from cDNA of dermal fibroblast 4 days post adenoviral infection. PCR was carried out in Platinum™ SuperFi II Green PCR Master Mix (Thermo Fisher Scientific, 12369010) with a PCR primer pair (Forward: 5’-AAACGCGTACAGGCTCATCT-3’; Reverse: 5’-TGGGAGAGGTCCTTGTCCAT-3’). The PCR product was purified by PCR purification kit (Zymo Research, D4013) and sent to Azenta Life Sciences for Sanger sequencing. PCR primers along with two additional primers that are more proximal to the G154R site (5’-CTGGCAGCACTCACCAACTA-3’; 5’-CAGCTGCGCCTCTTCATTG-3’) were used as sequencing primer to ensure successful sequencing with high quality. All primers used in the study were synthesized by Integrated DNA Technologies.

### 2.4 RT-PCR

cDNA of dermal fibroblast 4 days after adenoviral infection was used for quantification of TANGO2 expression. Real-time Taqman-PCR was assembled in qPCRBIO Probe Blue Mix (Genesee Scientific Corporation, 17-514) with 0.5μM of each primer (Forward: 5’-GGGAGCCTGATCCTATCGTT-3’; Reverse: 5’-TCCCAAAGCACAGCTTCC-3’) and 50nM Universal ProbeLibrary Probe #67 (Roche). RT-PCR was performed in QuantStudio™ 5 Real-Time PCR System (Thermo Fisher Scientific) and calculated using ^ΔΔ^Ct-method. Expression level of *GAPDH* (Forward: 5’-GCACAAGAGGAAGAGAGA GACC-3’; Reverse: 5’-AGGGGAGATTCAGTGTGGTG-3’; Probe #3) was used as internal control.

### 2.3 Seahorse Assay

Dermal fibroblasts of passage number 3-8 were dissociated by 0.25% Trypsin-EDTA (Thermo Fisher Scientific, 25200072), and seeded at 20K per well in the Seahorse XF96 cell culture microplates (Agilent Technologies, 102416100) in DMEM+10%FBS for maintenance until further usage. For fasting, fibroblasts culture was washed 3 times and the medium was replaced with a fasting medium composed of glucose-free DMEM (Thermo Fisher Scientific, A1443001) supplemented with 4.5g/L D-(+)-Galactose (Sigma, G0750) and 10% FBS.

iPSC-CMs were dissociated by 0.25% Trypsin-EDTA (Thermo Fisher Scientific, 25200072) or STEMdiff™ Cardiomyocyte Dissociation Kit (STEMCELL Technologies, 05025), and seeded at 50K per well in Matrigel-coated (Corning, 354277) Seahorse XF96 cell culture microplates in RPMI1640 supplemented with 1% B27, 10% KnockOut™ Serum Replacement (Thermo Fisher Scientific, 10828028), and 10 μM Y-27632 (MedChemExpress, HY-10071). Medium was replaced with RPMI1640+1% B27 two day after seeding for maintenance until further usage. For fasting, cardiomyocytes culture was washed 3 times and the medium was replaced with a fasting medium composed of glucose-free RPMI1640 (Thermo Fisher Scientific, 11879020) supplemented with 2g/L D-(+)-Galactose (Sigma, G0750) and 1% B27.

Oxygen consumption rate was measured in extracellular flux analyzer XFe96 (Agilent) with pyruvate (Sigma-Aldrich, P2256) or palmitate-BSA conjugate (Cayman Chemical, 29558) as substrate following the manufacturer’s protocol. 1.5 μM oligomycin (Sigma-Aldrich, 495455), 1 μM FCCP (Sigma-Aldrich, C2920), 5 μM rotenone (Sigma-Aldrich, 45656), and 5 μM antimycin A (Sigma-Aldrich, A8674) were used for the assay protocol (19). The maximum mitochondria respiratory capacity was measured as the difference between plateau value after injecting FCCP and the end-point value after injecting rotenone and antimycin A. OCR was normalized to DNA content in each well measured by CyQUANT® Cell Proliferation Assay kit (Thermo Fisher Scientific, C7026) (20).

## 3 RESULTS

### 3.1 Establishment of TANGO2 patient-derived dermal fibroblast and isogenic lines

We have previously derived and characterized an iPSC-CM from dermal fibroblasts of a TANGO2 patient carrying the recurrent homozygous ΔE3-9 mutation. Isogenic iPSC-CM lines were generated by infecting adenoviral vectors expressing WT-TANGO2, another recurring pathogenic mutation G154R or GFP control. G154R-TANGO2 is expected to be loss-of-function, which is equivalent to GFP from both clinical data and our in vitro results. We have shown that TANGO2 iPSC-CM expressing GFP or G154R-TANGO2 exhibits arrhythmias that resemble TANGO2 patients in crises, in contrast TANGO2 iPSC-CM expressing WT-TANGO2 fully corrected all the abnormalities in field potential and do not have arrhythmia (ref). These results support our use of this approach to generate isogenic cell lines to examine the effects of TANGO2 deficiency with high confidence.

We used a similar strategy to obtain isogenic dermal fibroblasts lines. The dermal fibroblasts from the same TANGO2 patient with homozygous ΔE3-9 mutation were used as the starting point (Figure 1A). Adenoviral vector infection of WT-TANGO2, G154R-TANGO2 and GFP control are used to generate isogenic fibroblasts lines. cDNA from fibroblasts of the ΔE3-9 patients and a WT control were used for genotyping (Figure 1B). As expected, there was no amplification from a primer pair detecting exon 7-8 of the human *TANGO2* transcript (ENST00000327374.9) from the ΔE3-9 cells. Primers targeting exon 1-2 showed amplifications from both WT control and ΔE3-9 cells, suggesting that the truncated mRNA which only contains the exon 1 and 2 was not degraded by nonsense mediated decay. However, the residue open reading frame contains only 19 amino acids, most are the mitochondrial targeting sequence, and are not expected to carry any TANGO2 specific function. Four days after adenoviral vector infection at MOI=1, the TANGO2 expression levels in ΔE3-9 fibroblast reached 7.8-fold (WT-TANGO2) and 6.6-fold (G154R-TANGO2) by comparing to four independent control fibroblast lines. As expected, GFP expressing fibroblasts do not express TANGO2 (Figure 1C). While the viral expressed TANGO2 levels are higher than those of the controls, they are still at a near physiological level. Genotypes of the isogenic lines were further confirmed by Sanger sequencing (Figure 1D).

**FIGURE 1.**
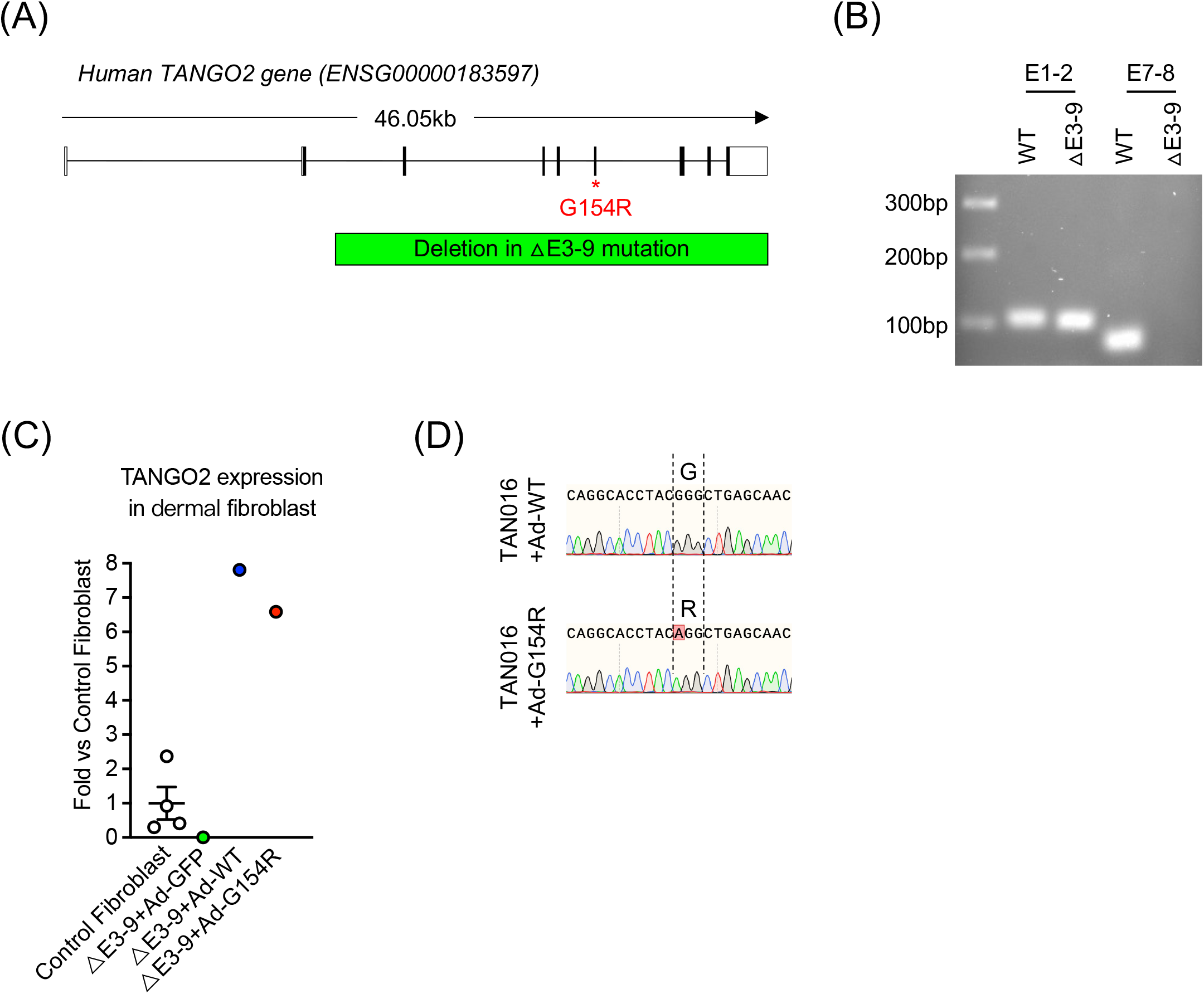
Generation of △E3-9 dermal fibroblast and its isogenic lines by adenoviral expression. A, Location of G154R and △E3-9 (deletion of Exon 3-9) mutations in human *TANGO2* gene. Box indicates exons and line indicates introns. Coding region in exons is colored in black. B, Genotyping of △E3-9 fibroblast using primer pairs targeting exon 1-2 (E1-2) or exon 7-8 (E7-8) of WT-TANGO2 transcript. C-D. A WT fibroblast was used as positive control. C, TANGO2 expression level in △E3-9 fibroblast, △E3-9 fibroblast with adenoviral expression of WT-TANGO2, G154R-TANGO2, and GFP control, and 4 independent control fibroblast lines from healthy donors. Samples were harvested 4 days post-infection. GAPDH was used as internal control. Data were normalized to the level of control fibroblasts. D, Sanger sequencing showing desired genotype in △E3-9 fibroblast with adenoviral expression of WT-TANGO2 and G154R-TANGO2. PCR product amplified from cDNA was used for sequencing.

### 3.2 Mitochondrial OXPHOS function is normal in TANGO2 deficient dermal fibroblasts

The mitochondrial OXPHOS function of isogenic fibroblast lines were interrogated by the Seahorse Cell Mito Stress Assay. We first started with two TANGO2 deficient fibroblast lines that carry the ΔE3-9 or G154R mutation, along with a set of independent controls dermal fibroblast lines from healthy donors (Figure 2 and Supplementary Figure 1). Before the assay, cells were subjected to two culture conditions. First, a baseline condition where cells were maintained in regular glucose containing medium. Second, a fasting condition where cells were maintained in a fasting medium which was made by replacing the glucose with the same amount of galactose in the regular medium. Because galactose does not generate net ATP via glycolytic metabolism, the galactose medium forces the cellular energy production to depend more on OXPHOS, which mimics the nutrient stress induced by glucose deprivation or fasting (21–23). In addition, fasting can increase PDK4 (pyruvate dehydrogenase kinase isoenzyme 4) expression which suppresses glycolytic pathway but activates fatty acid oxidation (24). Consistently, we found 4-day fasting greatly reduced the baseline and maximal pyruvate-dependent OCR (p<0.05, two-way ANOVA) with a trend of increase in maximal palmitate-dependent OCR in both control and TANGO2 deficient groups (Figure 2), suggesting successful induction of fasting effect by galactose medium. However, there was no significant difference in pyruvate or palmitate-dependent OCR between the TANGO2 deficient (ΔE3-9+GFP or ΔE3-9+G154R) and 6 independent control (WT) lines under baseline or fasting conditions (Figure 2). When comparing each line individually, we found the OCRs of two TANGO2 deficient lines lie within the variation of the OCRs of the healthy control set (Supplementary Figure 1). Thus, this data indicate TANGO2 deficiency may not significantly impair mitochondrial OXPHOS functions.

**FIGURE 2.**
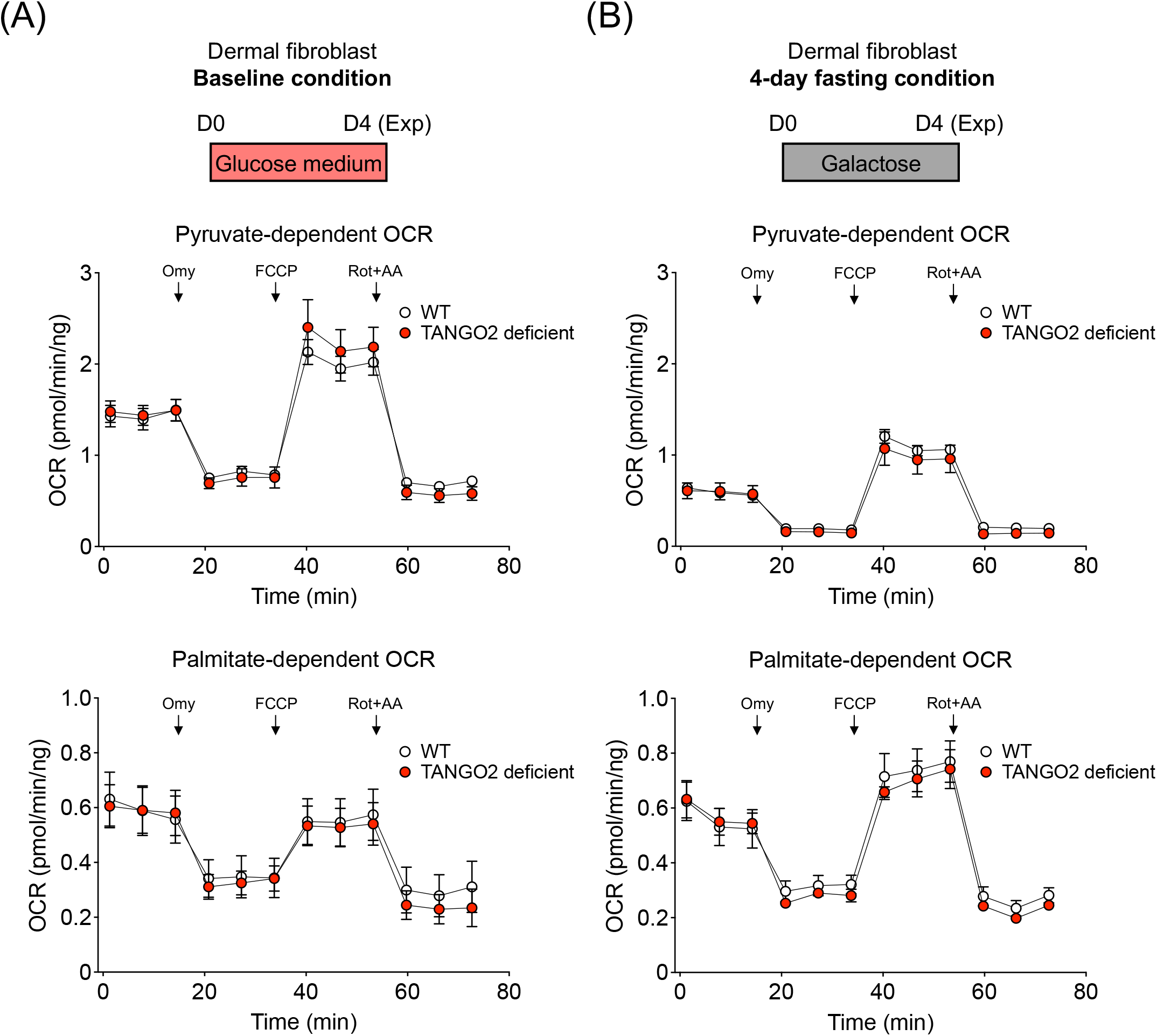
TANGO2 deficient dermal fibroblasts have normal OXPHOS level under 4-day glucose (baseline) and 4-day galactose (fasting) condition. A, Measurement of oxygen consumption rate (ORC) in TANGO2 deficient and WT fibroblast lines under baseline condition where cells were maintained in glucose medium for 1 day before the Seahorse experiment (Exp). Data in TANGO2 deficient group is calculated as the averaged data from two fibroblast lines that carry △E3-9 and G154R mutations. Data in WT groups are averaged data from 6 dermal fibroblast lines from healthy donors. B, ORC measurement in TANGO2 deficient and WT fibroblast lines under a fasting condition where cells were maintained in galactose medium for 4 day before the Seahorse experiment (Exp). Pyruvate (Upper panel) or palmitate (lower panel) was used substrate in the Seahorse assay. Arrows indicate injection of oligomycin (Omy), FCCP, Rotenone (Rot) and antimycin A (AA). N=2 for TANGO2 deficient group and N=5-6 for WT group. Data are presented as mean ± SEM. Error bars smaller than the circles are not shown.

The large variation in healthy controls underscores the importance of using isogenic lines. Thus, we implemented the isogenic fibroblast lines with adenoviral infection (Figure 1) to validate the results. Following a similar experimental design, the isogenic fibroblasts were subjected to two culture conditions:

1. A baseline condition where fibroblasts were maintained in regular glucose containing medium for 7 days after adenoviral infection, which confers sufficient time for the protein to be produced (Figure 3A).
2. A fasting condition where cells were maintained in regular medium for 3 days followed by a 4-day challenge in fasting medium (Figure 3B).

**FIGURE 3.**
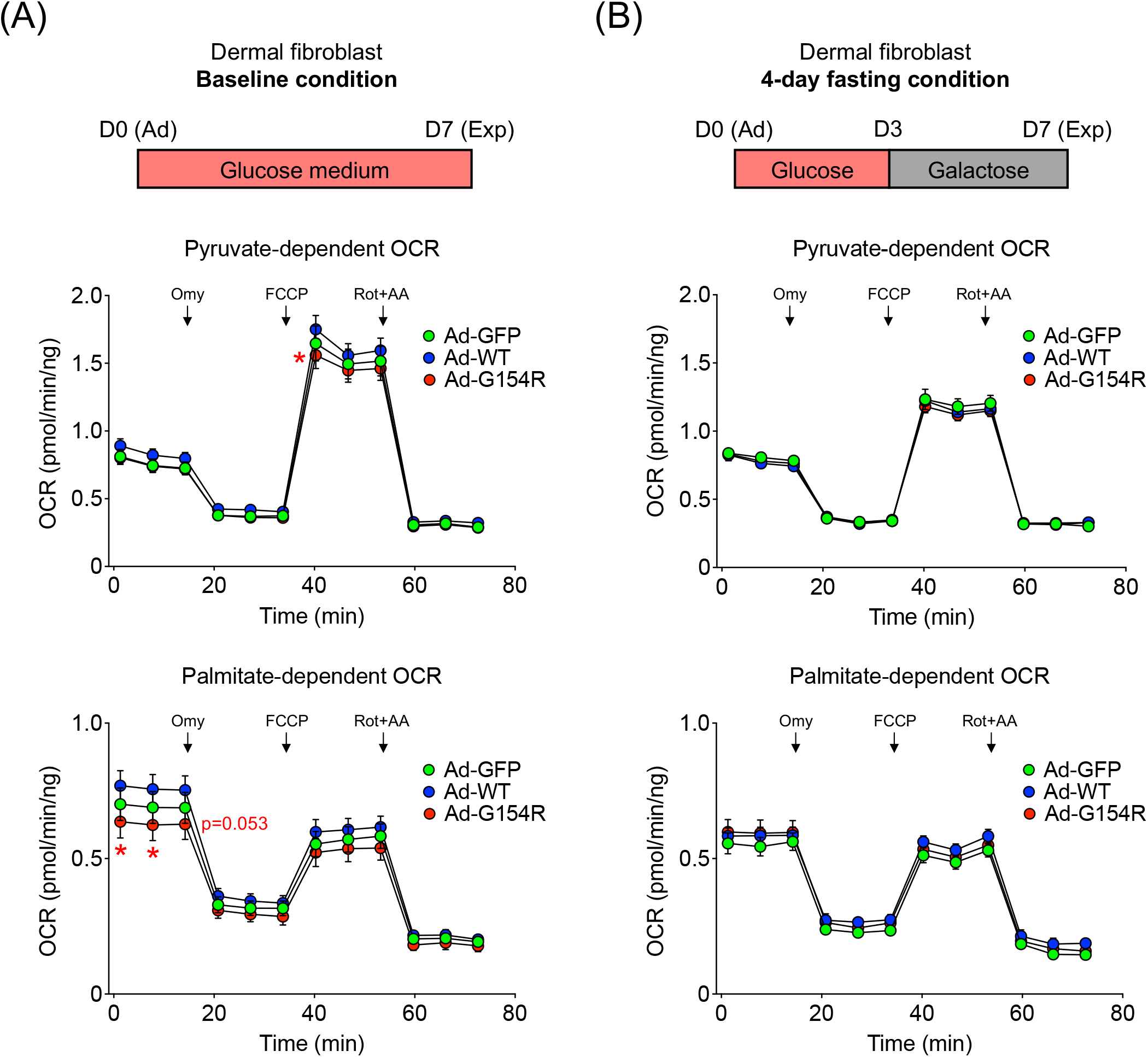
Isogenic TANGO2 deficient dermal fibroblast had normal OXPHOS function under 7-day glucose (baseline) condition or 3-day glucose+4-day galactose (fasting) condition. A, Measurement of oxygen consumption rate (ORC) in △E3-9 TANGO2 dermal fibroblast with adenoviral expression of WT-TANGO2, G154R-TANGO2, or GFP control under baseline condition. B, OCR measurement in △E3-9 TANGO2 dermal fibroblast with adenoviral expression of WT-TANGO2, G154R-TANGO2, or GFP control under fasting condition. Adenoviral infection (Ad) and Seahorse experiment (Exp) were carried out on D0 and D7, respectively. Pyruvate (Upper panel) or palmitate (lower panel) was used substrate in the Seahorse assay. Arrows indicate injection of oligomycin (Omy), FCCP, Rotenone (Rot) and antimycin A (AA). N=14-16. *p<0.05 vs Ad-WT. Statistical difference is determined by two-way ANOVA. Multiple comparison is corrected by Tukey method with α=0.05. Data are presented as mean ± SEM. Error bars smaller than the circles are not shown.

When using pyruvate as substrate, we observed a slightly higher (12.22%) maximal OCR in WT group than that in the G154R group at baseline condition (Figure 3A upper panel). As expected, fasting reduced the maximal OCR in all three isogenic fibroblast lines compared to that at baseline (p<0.05, two-way ANOVA), however, when comparing among groups no difference was found (Figure 3B upper panel). When using palmitate as the substrate, there’s a trend of increase in basal OCR of WT group vs G154R group under baseline condition (Figure 3A lower panel), while no difference in OCR was found under fasting condition (Figure 3B lower panel). Notably, GFP group had similar level of OCR to that of WT under any of the experimental conditions (Figure 3). Therefore, results from isogenic control fibroblasts largely agree with the results from independent fibroblast lines (Figure 2), suggesting normal mitochondrial OXPHOS function in TANGO2 deficient dermal fibroblast.

### 3.3 Mitochondrial OXPHOS function is normal in TANGO2 deficient iPSC-CM

Because of the energy-demanding nature of heart and the severe cardiac phenotype exhibited in TANGO2 patients during crises, one may argue that the mitochondrial function in cardiomyocytes is more susceptible to TANGO2 deficiency than that in dermal fibroblast. To address this possibility, we used our isogenic iPSC-CM models to interrogate the effects of TANGO2 deficiency on mitochondrial OXPHOS function. For baseline condition, short term of 4 days (Figure 4A), medium term of 7 days (Figure 5A), and long term of 14 days (Figure 6A) under glucose medium culture were performed to ensure the time required to reach sufficient level of ectopic expression and to observe significant alteration in mitochondrial function. For the fasting condition, short-term fasting for 1 day (Figure 4B) and long-term fasting for 4 days (Figure 5B and 6B) were performed. Fasting longer than 4 days was found to cause significant cytopathic effects and detachment of iPSC-CM in all groups.

**FIGURE 4.**
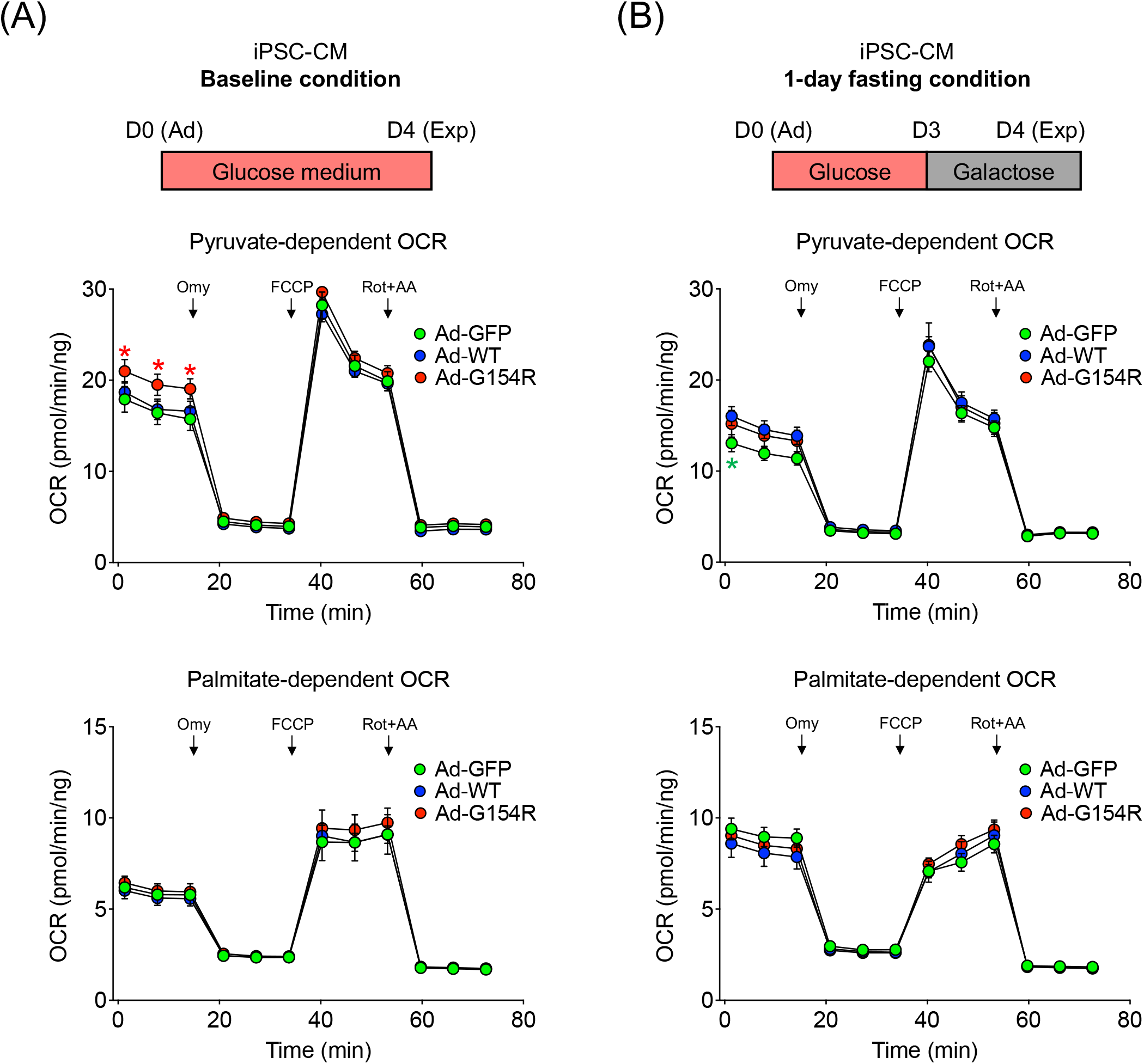
Isogenic TANGO2 deficient iPSC-CM had normal OXPHOS function under 4-day glucose (baseline) condition or 3-day glucose+1-day galactose (fasting) condition. A, Measurement of oxygen consumption rate (ORC) in △E3-9 TANGO2 iPSC-CM with adenoviral expression of WT-TANGO2, G154R-TANGO2, or GFP control under baseline condition where cells were maintained in glucose medium for 4 days. B, ORC measurement in △E3-9 TANGO2 dermal fibroblast with adenoviral expression of WT-TANGO2, G154R-TANGO2, or GFP control under a fasting condition where cells were maintained in glucose medium for 3 days followed by a 1-day culture in galactose medium. Adenoviral infection (Ad) and Seahorse experiment (Exp) were carried out on D0 and D4, respectively. Pyruvate (Upper panel) or palmitate (lower panel) was used substrate in the Seahorse assay. Arrows indicate injection of oligomycin (Omy), FCCP, Rotenone (Rot) and antimycin A (AA). N=7-8. *p<0.05 vs Ad-WT. Statistical difference is determined by two-way ANOVA. Multiple comparison is corrected by Tukey method with α=0.05. Data are presented as mean ± SEM. Error bars smaller than the circles are not shown.

**FIGURE 5.**
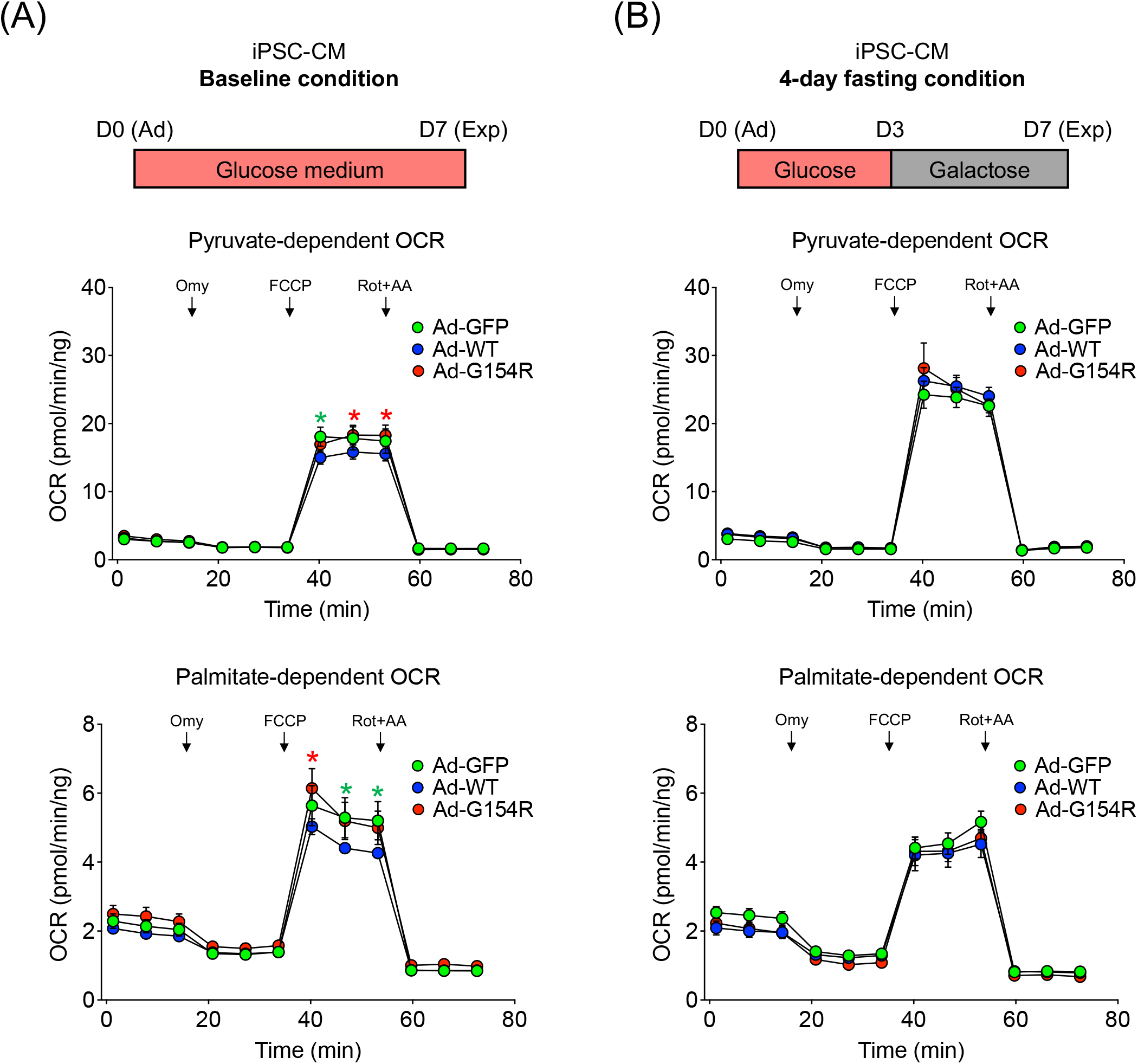
Isogenic TANGO2 deficient iPSC-CM had normal OXPHOS function under 7-day glucose (baseline) condition or 3-day glucose+4-day galactose (fasting) condition. A,Measurement of oxygen consumption rate (ORC) in △E3-9 TANGO2 iPSC-CM with adenoviralexpression of WT-TANGO2, G154R-TANGO2, or GFP control under baseline condition where cells were maintained in glucose medium for 7 days. B, ORC measurement in △E3-9 TANGO2 dermal fibroblast with adenoviral expression of WT-TANGO2, G154R-TANGO2, or GFP control under a fasting condition where cells were maintained in glucose medium for 3 days followed by a 4-day culture in galactose medium. Adenoviral infection (Ad) and Seahorse experiment (Exp) were carried out on D0 and D4, respectively. Pyruvate (Upper panel) or palmitate (lower panel) was used substrate in the Seahorse assay. Arrows indicate injection of oligomycin (Omy), FCCP, Rotenone (Rot) and antimycin A (AA). N=7-8. *p<0.05 vs Ad-WT. Statistical difference is determined by two-way ANOVA. Multiple comparison is corrected by Tukey method with α=0.05. Data are presented as mean ± SEM. Error bars smaller than the circles are not shown.

**FIGURE 6.**
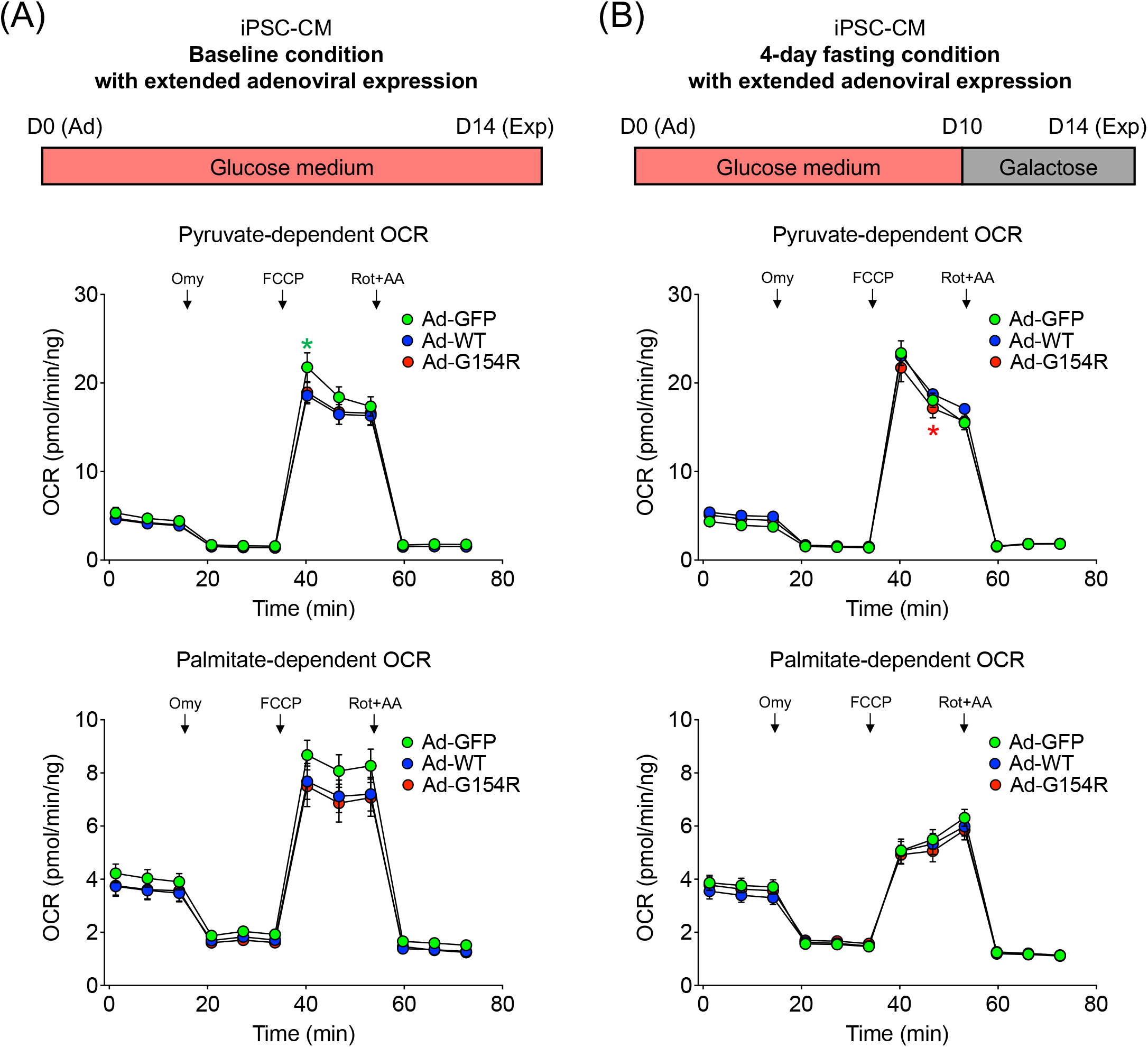
Isogenic TANGO2 deficient iPSC-CM had normal OXPHOS function under 14-day glucose (baseline) condition or 10-day glucose+4-day galactose (fasting) condition. A, Measurement of oxygen consumption rate (ORC) in △E3-9 TANGO2 iPSC-CM with adenoviral expression of WT-TANGO2, G154R-TANGO2, or GFP control under baseline condition where cells were maintained in glucose medium for 14 days. B, ORC measurement in △E3-9 TANGO2 dermal fibroblast with adenoviral expression of WT-TANGO2, G154R-TANGO2, or GFP control under a fasting condition where cells were maintained in glucose medium for 10 days followed by a 4-day culture in galactose medium. Adenoviral infection (Ad) and Seahorse experiment (Exp) were carried out on D0 and D14, respectively. Pyruvate (Upper panel) or palmitate (lower panel) was used substrate in the Seahorse assay. Arrows indicate injection of oligomycin (Omy), FCCP, Rotenone (Rot) and antimycin A (AA). N=7-8. *p<0.05 vs Ad-WT. Statistical difference is determined by two-way ANOVA. Multiple comparison is corrected by Tukey method with α=0.05. Data are presented as mean ± SEM. Error bars smaller than the circles are not shown.

Consistent with the fasting effects observed in dermal fibroblast, 1-day fasting significantly reduced the maximal pyruvate-dependent OCR (p<0.05, two-way ANOVA) but increased basal palmitate-dependent OCR (p<0.05, two-way ANOVA) in all three isogenic iPSC-CM lines (Figure 4). When comparing GFP/G154R group to WT group, we found normal or even greater OCR in GFP or G154R group (Figure 4), except a slight decrease (22.5%) in basal pyruvate-dependent OCR in GFP group after 1-day fasting. (Figure 4B upper panel). Data from longer time points also showed that OCR levels of GFP/G154R iPSC-CM and WT iPSC-CM are comparable or even higher (Figure 5–6), except a subtle (9.37%) decrease in maximal pyruvate-dependent OCR in G154R group under the 10-day glucose+4-day fasting condition (Figure 6B upper panel). Taken together, our results suggest that TANGO2 deficient cardiomyocytes have normal mitochondrial OXPHOS function, and response to fasting. The primary defect that leads to the metabolic crisis and lethal cardiac arrythmias in TANGO2 patients are not related to an impaired OXPHOS function in the cardiomyocytes.

## 4 DISCUSSIONS

TANGO2-related disease was first described in 2016 (3). Deciphering the disease etiology has been challenging largely due to lack of basic understanding of TANGO function in human. *TANGO2* was first identified through a genome-wide RNA inference screening for genes involved in protein secretion and Golgi organization in a *Drosophila* cell line in 2006 (25). Nevertheless, sequence alignment between human (Q6ICL3 in UniPort) and *Drosophila* protein orthologs (Q9VYA8 in Uniport) revealed only 27.97% identity, suggesting TANGO2 may have different function in human. Increasing attention has been drawn to the association of TANGO2 with mitochondrial function in the field, supported by the similarity between clinical feature of TANGO2-related disease and inborn errors of metabolism (1–14), as well as the prediction of a mitochondrial targeting sequence at the N-terminus (2). So far there has been only 4 studies investigating the TANGO2 mitochondrial localization and the impact of TANGO2 deficiency on mitochondrial respiratory function, however, results are inconsistent from different studies (2, 5, 13, 14). The inconsistency likely represents the large variation among the patient-derived dermal fibroblasts, as is demonstrated in our study as well (Supplementary Figure 1). Therefore, it is difficult to obtain convincing and conclusive data without a sizable number of fibroblast cell lines from independent control and disease subjects. Moreover, as metabolic crises-induced cardiac arrhythmias are the leading cause of mortality in TANGO2 patients, study utilizing cardiac models can provide more relevant information on the pathogenesis of TANGO2-associated arrhythmias, which has been lacking in the field.

To overcome these limitations in previous studies, we implemented both isogenic dermal fibroblast lines and isogenic iPSC-CM lines aiming to directly answer whether TANGO2 deficiency leads to primary defects in mitochondrial OXPHOS function. The isogenic fibroblast or iPSC-CM lines generated by adenoviral expression from the TANGO2 deficient cells (ΔE3-9) have the identical genetic background, thus eliminating the variation from different genetic background. Further, as the mutations were introduced post differentiation using viral vectors, the variation associated with cardiomyocytes differentiation can also be minimized. Our models provide the ideal experimental setting to interrogate cellular function between TANGO2 deficiency and control groups with high confidence in the most relevant human cell type.

As fasting is a known trigger for metabolic crisis in TANGO2 patients (1, 7), we also challenged the cells with fasting medium where glucose was replaced with galactose in the culture medium for 1 or 4 days. Expected fasting effects including suppression of glycolytic pathway and elevation in fatty acid oxidation level (24) were observed in both fibroblast and iPSC-CM models (Figure 2–4 and Supplementary Figure 1). The mitochondrial function was examined using Mito Stress Test with either pyruvate or palmitate as the substrate to measure oxygen consumption rate (OCR) in Seahorse assay, a widely used method to measure the key parameters of mitochondrial respiration (26, 27). Our data showed that under most of the conditions that were tested in this study both TANGO2 deficient dermal fibroblast and iPSC-CM have normal or even higher level of mitochondrial OXPHOS compared to WT control (Figure 2–6 and Supplementary Figure 1). It is important to note that while there are a few isolated conditions that TANGO2 deficient cells exhibited minor reduction in OCR levels (Figure 3A, Figure 4B upper panel, and Figure 6B upper panel), none were exaggerated by fasting or prolonged fasting or consistent between the 2 TANGO2 deficient genotypes (GFP and G154R). Thus, they are more likely isolated experimental variations without significant biological implications. Therefore, we conclude that TANGO2 deficiency is unlikely to cause a primary defect in mitochondrial OXPHOS function which leads to lethal arrythmias in TANGO2 patients.

Our results do not rule out the possibility that indirect effects on mitochondrial OXPHOS function may occur during metabolic crisis in TANGO2 patients in vivo as is suggested by a previous publication (5). Animal model with TANGO2 deficiency is required to examine possible influence from other organ system on heart. In addition, although our conclusion was primarily based on the results from iPSC-CM model, it is possible that TANGO2 deficiency may impair metabolic OXPHOS function in other type of cells, which may contribute to the phenotype seen in other organs such as neurological defects. Future study using iPSC-derived neurons or other type of cells may address this hypothesis. Finally, it remains an open question whether other OXPHOS-unrelated mitochondrial function may be impaired by TANGO2 deficiency, for example, ER-mitochondrial contacts (28) which may be associated with the recently proposed dual function of TANGO2 in mitochondrial and ER/Golgi network (14). Mitochondrial ROS level will also need to be investigated in future study, although dramatic change is not expected as otherwise impaired OXPHOS function should have been observed (29, 30).

## ACKNOWLEDGEMENTS

We thank the Mouse Metabolism and Phenotyping Core at Baylor College of Medicine for assisting the Seahorse study. We thank Undiagnosed Diseases Network (UND) for providing resource and access to human dermal fibroblasts.

## AUTHOR CONTRIBUTIONS

Seema R. Lalani, Christina Y. Miyake, and Lilei Zhang conceived the project. Weiyi Xu, and Lilei Zhang designed the study. Weiyi Xu, Yingqiong Cao, Lorren Cantú, and Eleni Nasiotis performed the experiments. Weiyi Xu, Lorren Cantú, and Eleni Nasiotis performed data analysis. Weiyi Xu, Seema R. Lalani, Christina Y. Miyake, and Lilei Zhang wrote the manuscript.

## ETHICS STATEMENT

The use of patient-derived fibroblasts was approved by the Institutional Review Board of Baylor College of Medicine (H-43240). Written informed consents from subjects were obtained by the study team.

---

**Supplementary Figure 1.**
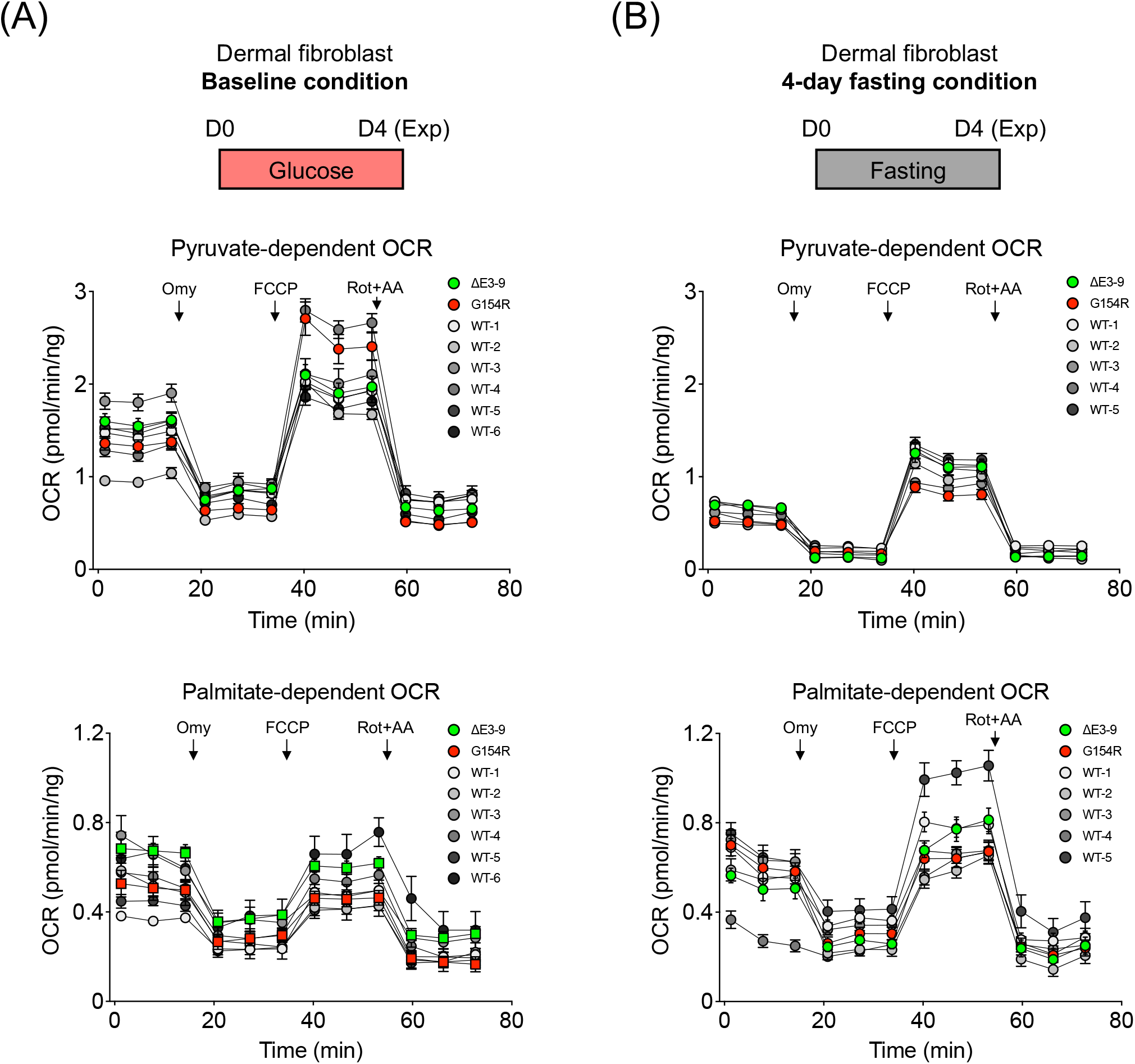
Mitochondrial OXPHOS capacity of two TANGO2 deficient dermal fibroblasts lines lies within the variation of that of WT dermal fibroblast lines under 4-day glucose (baseline) and 4-day galactose (fasting) condition. A, Measurement of oxygen consumption rate (ORC) in △E3-9 TANGO2, G154R-TANGO2, and 6 WT dermal fibroblast lines from healthy donors under baseline condition where cells were maintained in glucose medium for 1 day before the experiment (Exp). B, ORC measurement in △E3-9 TANGO2, G154R-TANGO2, and 6 WT dermal fibroblast lines from healthy donors under a fasting condition where cells were maintained in galactose medium for 4 day before the experiment (Exp). Pyruvate (Upper panel) or palmitate (lower panel) was used substrate in the Seahorse assay. Arrows indicate injection of oligomycin (Omy), FCCP, Rotenone (Rot) and antimycin A (AA). N=5-12. Data are presented as mean ± SEM. Error bars smaller than the circles are not shown.

